# Structural characterization of MAM01 and other cost-effective engineered monoclonal antibodies for malaria prevention

**DOI:** 10.1101/2025.08.24.672003

**Authors:** Monika Jain, Sashank Agrawal, Gonzalo E. González-Páez, Re’em Moskovitz, Randal R. Ketchem, Katherine L. Williams, Daniel E. Emerling, Ian A. Wilson

## Abstract

Long-lasting and effective vaccines or monoclonal antibodies (mAbs) for malaria prevention are highly beneficial for people of all ages living in malaria-endemic regions. Recently, two highly protective human mAbs AB000224 (IGHV3-49/IGLV1-40) and AB007088 (IGHV3-33/IGKV1-5), which are encoded by different germline genes, were re-engineered and renamed as MAM01 and MS-1805, respectively, to improve their half-life, developability (including manufacturing), and cost-effectiveness according to WHO guidelines. MAM01 has completed phase 1 and 1b clinical trials (safety / PK / challenge) in US healthy naive adults and is in phase 1 trials (age-de-escalation studies) in Uganda. Here, we determined crystal structures of the antigen-binding fragments (Fabs) of engineered MAM01, MS-1805, and 7088 in complex with different regions of the *Plasmodium falciparum* (Pf) circumsporozoite protein (CSP), including junctional, minor, and major repeat regions. Notably, Fab 7088 features an extended CDRL3 comprising 10 amino acids (CDRL3:10) instead of the typical 8 amino acid CDRL3 (CDRL3:8) for V_H_3-33/V_K_1-5-encoded mAbs, revealing unique folding within its germline context. Structural comparisons showed that engineered antibodies retain the key molecular and hydrogen bond interactions with no significant conformational changes or loss in binding affinity. These findings demonstrate that the efficacy and affordability of human mAbs can be enhanced by selectively mutating residues in antibody framework regions without compromising binding affinity or epitope interaction.

## Introduction

Malaria is still a major global concern and public health threat due to emerging environmental factors associated with climate change and vector resistance to insecticides (1, 2). In 2023, 263 million cases of malaria were reported worldwide, and infection with *Plasmodium falciparum*, the deadliest species of the malaria parasite, led to high mortality among young children and pregnant women, particularly in sub-Saharan Africa and Southeast Asia (3). The *P. falciparum* circumsporozoite protein (PfCSP) densely covers the sporozoite surface and plays a critical role in sporozoite development from the mosquito midgut to human hepatocyte invasion (4). PfCSP (Pf reference strain 3D7) comprises an N-terminal domain, a junctional region (with one NPDP tetrapeptide repeat), a central minor repeat region (4 NVDP repeats), a major repeat region (38 NANP repeats) and a C-terminal α-thrombospondin repeat (αTSR) domain (5).

Recently, RTS,S/AS01(Mosquirix) and R21/Matrix-M have become the two most effective vaccines approved by WHO for children living in malaria-endemic regions (6). These subunit vaccines contain a truncated form of PfCSP, comprising only 19 of the NANP major repeats and the entire C-terminal region, but were reported to have limitations in their efficacy and duration of the immune response (7-9). However, several highly protective human monoclonal antibodies (mAb) have been isolated from participants vaccinated with RTS,S/AS01, including mAbs 317 and 311 (10), or immunized with an attenuated whole sporozoite vaccine (PfSPZ vaccine), including mAb L9 and CIS43 (11, 12). These mAbs aim to block the initial stages of infection by targeting conserved regions on the sporozoite. Treatment with these mAbs, which target different regions of the CSP, inhibit sporozoite infection and demonstrate protection from malaria in mouse models (13-15). Recent clinical trials have shown that human L9 and CIS43 mAbs, which target the CSP NVDP minor region and NPDP junctional region, respectively, prevent infection against malaria (16-18). Both antibodies were further optimized by introducing LS point mutations (M428L/N434S) in the fragment crystallizable (Fc) region C_H_3 domain to extend their half-life and were named L9LS and CIS43LS (17, 19).

The mAbs AB-000224 and AB-007088, like 317 and 311, were also isolated from the circulating B cells of protected individuals administered with the RTS,S/AS01 vaccine, and showed superior protection and high sporozoite inhibition in the liver compared to the highly efficacious mAb AB-000317 (20-22). Antibodies AB-000224 and AB-007088 were subsequently engineered by introducing specific mutations in the framework regions of their variable domains along with LS mutations in the Fc region to enhance their stability and extend half-life for use as therapeutics (23, 24). The engineered AB-000224, renamed MS-1797 (MAM01), was selected as the main drug candidate, and advanced into clinical development, whereas the engineered AB-007088, renamed MS-1805, was designated as a backup candidate for malaria prevention (22, 25).

MAM01 and MS-1805 retain their binding to peptides derived from different CSP regions, similar to the antibodies AB000224 and AB007088. The modifications in these antibodies were characterized by pharmacological and biophysical assays to assess optimized manufacturability and long-acting formulation. However, it is crucial to also assess the structural and conformational stability of these engineered mAbs. Here, we report X-ray structures of engineered MAM01 Fab and 7088 Fab, along with its engineered counterpart MS-1805 in complex with different CSP-derived peptides. We observed that 7088 Fab possesses an extended CDRL3 of 10 amino acids (CDRL3:10), which differs from the typical CDRL3 of 8 amino acids (CDRL3:8) observed in other heavy chain IGHV3-33 associated with light chain IGKV1-5 germline gene (26, 27). With the aid of crystal structures of Fab 7088 and MS-1805, we observed how the long CDRL3 of 10 amino acids interacts with the antigen and how the long CDRL3 is accommodated in the context of the IGHV3-33/IGKV1-5 germline-encoded antibody.

Moreover, structural characterization of the engineered antibodies MAM01 and MS-1805 showed no significant conformational changes in the overall antibody structure, in their complementarity-determining regions (CDRs), or in their hydrogen bond interactions, compared to the antibodies AB000224 and AB007088. These findings confirm that specific modifications can be applied to the naturally occurring antibodies to improve their stability, manufacturability, and cost-effectiveness.

## Results

### Binding affinity of Fabs 224 and 7088 and engineered Fabs MAM01 and MS-1805 to PfCSP-derived epitopes of different regions

The antibody sequences of AB000224 and AB007088 were identified from plasmablasts isolated from protected participants enrolled in a phase 2a clinical trial of RTS,S/AS01 (21). For clinical trials, these mAbs were engineered by mutating specific residues and renamed MAM01 and MS-1805, respectively, which enhanced their stability, half-life, and manufacturability at lower cost (22). For MAM01, mutations were introduced in the variable domain framework region of the heavy chain of AB000224 antibody. Proline, isoleucine, and threonine residues were replaced with serine at amino acid position 21, threonine at position 77, and alanine at position 84 and 93 (Kabat numbering) (Fig. 1A). Additionally, in the Fv region of the light chain, glutamic acid and arginine residues were replaced with glutamine at position 1 and threonine at position 42, respectively (Fig. 1A). In contrast, modifications in engineered AB007088 were restricted to the heavy chain with no changes in the light chain. For MS-1805, specific mutations included replacing alanine at position 28 and isoleucine at position 68 and 110 with threonine, while glycine at 82B, serine at position 79 and threonine at position 40 were replaced with serine, tyrosine and alanine respectively (Fig. 1B). For both antibodies, LS mutations at (M428L/N434S) were introduced in the Fc region to extend half-life, but the effect of these mutations are not analyzed here as we are focusing on the Fabs of these antibodies.

**Figure 1.**
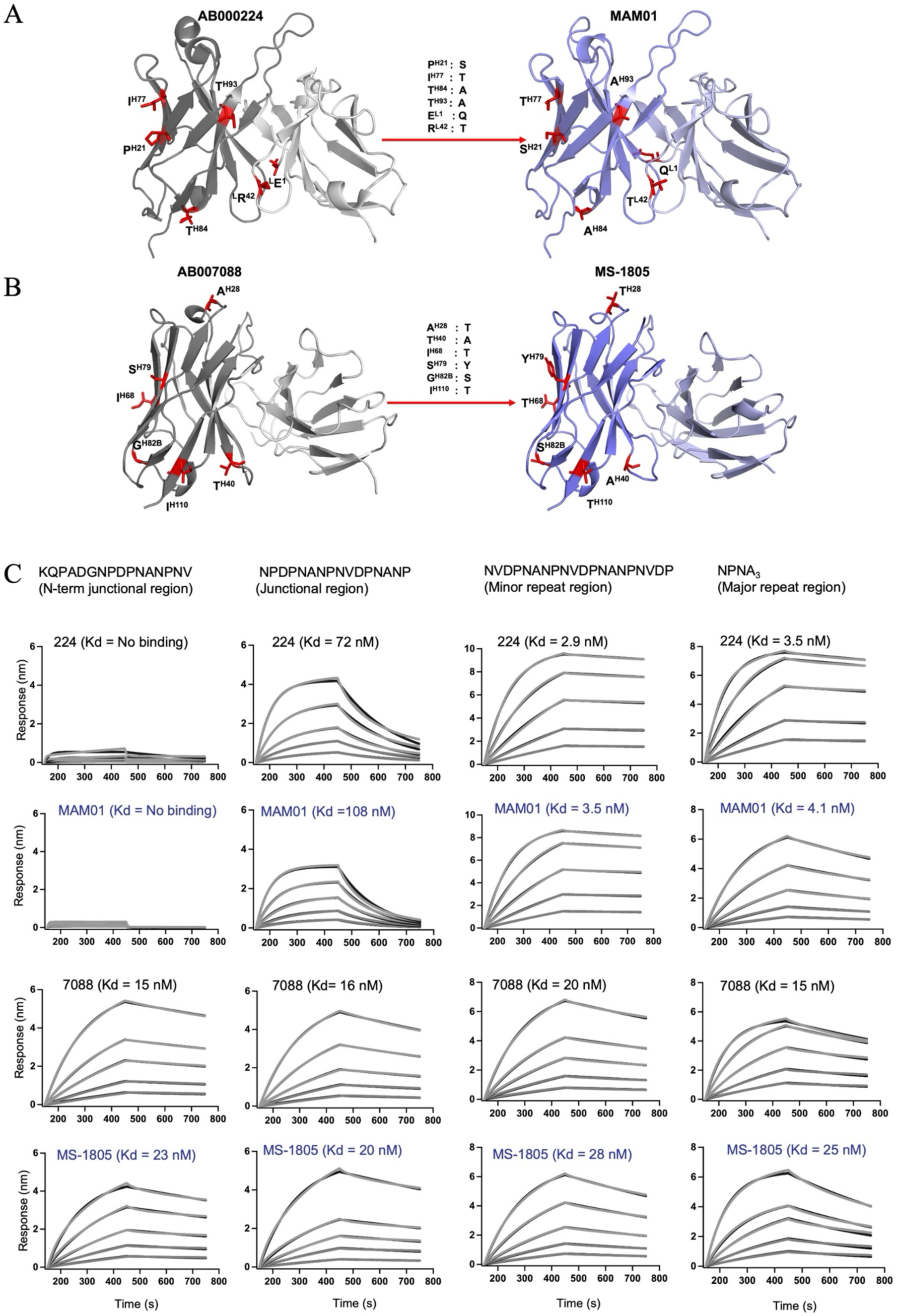
Binding kinetics for Fabs with PfCSP-derived epitopes from different regions analyzed in this study. A-B) Cartoon representation of the polypeptide backbone of the heavy and light variable chains for Fabs 224 and 7088 and engineered Fabs MAM01and MS-1805. For 224 and 7088 Fabs, the heavy chain is shown in dark grey and light chain in light grey, while for engineered Fabs, the heavy chain is in dark blue and light chain in light blue. Mutations introduced primarily in the framework regions of Fabs 224 and 7088 to generate MAM01 and MS-1805, respectively, are marked in red and the residues labeled with their locations in superscript (Kabat numbering). C) Bio-layer interferometry analysis of binding kinetics for both 224 and 7088 and their engineered Fabs with various PfCSP-derived peptides. Binding curves and fitted curves (1:1 global fitting model) are shown in grey and black, respectively. Serial dilutions of Fab concentrations are 200, 100, 50, 25, 12.5 nM from top to bottom and K_d_ values of each Fabs to their respective peptides are shown. (See also Table S1).

We analyzed and compared the binding affinity of Fabs 224 and 7088 with the engineered Fab MAM01 and MS-1805 for CSP-derived peptides from different regions, including the N-terminal junctional region (KQPADGNPDPNANPNV), junctional region (NPDPNANPNVDPNANP), minor repeat region (NVDPNANPNVDPNANPNVDP), and major repeat region (NPNA_3_) (Fig. 1C). Bio-layer interferometry (BLI) was performed to analyze the binding affinity and kinetics of these Fabs to the CSP-derived peptides (Table S1). We observed that Fab MAM01 binds with high affinity (K_D_ 3-4 nM) to the minor and major repeat regions compared to the junctional region (K_D_ 108 nM), retaining similar binding affinity to these peptides as Fab 224 (Fig. 1C). However, Fab MS-1805 binds to all of these peptides with high affinity (K_D_ 20-28 nM), which were very similar to Fab 7088 (K_D_ 15-20 nM). Notably, we found that Fab 7088 and MS-1805 also bind to peptides from the N-terminal junctional region (KQPADG), while Fab 224 and MAM01 showed no binding to this region. These results suggest that the N-terminus of the CSP junctional region of PfCSP might be involved in binding to Fabs 7088 and MS-1805 (Fig. 1C).

### Crystal structures of engineered MAM01 Fab with CSP-derived peptides of junctional, minor repeat, short major repeat and long major repeat regions

Crystal structures of MAM01 Fab in complex with CSP-derived peptides from the junctional region, minor repeat region, short major repeat region (NPNA_3_), and long major repeat region (NANP_6_) were determined to elucidate the structural basis of MAM01 binding to various CSP-derived peptides (Fig. 2A). The different crystals diffracted to 2.71 Å, 1.84 Å, 1.54 Å and 1.48 Å resolution, respectively (Table S2).

**Figure 2.**
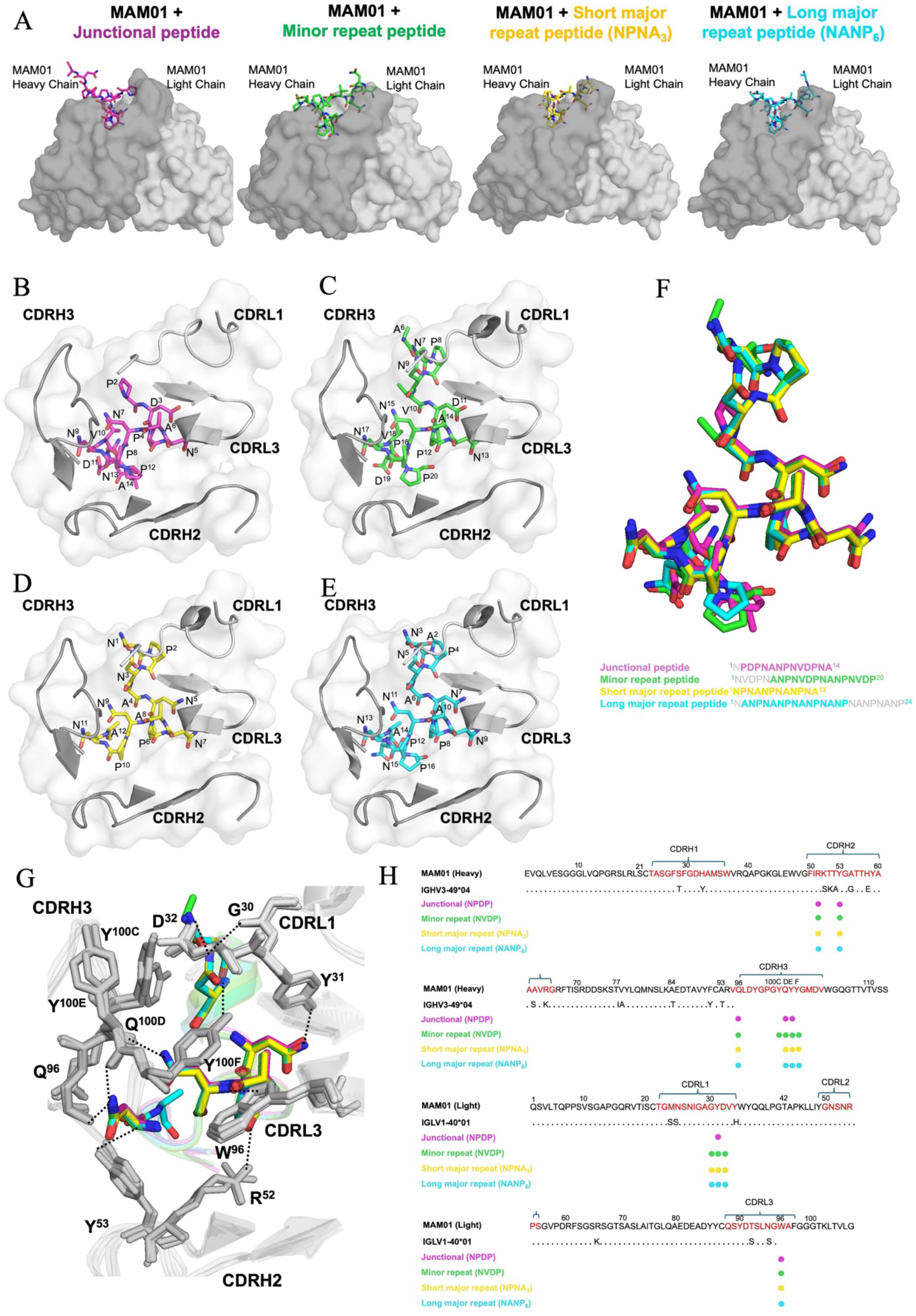
Crystal structures of MAM01 Fab with CSP-derived peptides. A) Surface representation of MAM01 Fab bound to peptides from junctional (magenta), minor repeat (green), short major repeat (yellow), and long major repeat (cyan) regions. Variable domains are colored dark grey for heavy chain and light grey for light chain. B-E) Close-up view of MAM01 CDRs interacting with each peptide: B) junctional (magenta), C) minor repeat (green), D) short major repeat (yellow), and E) long major repeat (cyan). F) Superimposed peptides are colored according to sequence. G) Hydrogen-bonding interactions between MAM01 and peptides, are highlighted in black dashes. H) MAM01 Fab sequence (Kabat numbering) with CDR residues interacting with each peptide marked as closed circles (colored by peptide region).

The interactions between Fab MAM01 and the peptides are mediated through heavy chain CDRH2 and CDRH3, and light chain CDRL1 and CDRL3. In the MAM01 Fab complex with junctional peptide (^1^NPDPNANPNVDPNA^14^), residues were assigned numbers from 1 to 14, although the electron density was not well defined for Asn^1^ (Fig. 2B). Similarly, for minor repeat peptide (^1^NVDPNANPNVDPNANPNVDP^20^), residues were numbered from 1 to 20, but no electron density was visible for Asn^1^ to Asn^5^ (Fig. 2C). In the MAM01 complex with short major repeat (^1^NPNANPNANPNA^12^), residues were numbered from 1 to 12 with well-defined electron density for all residues (Fig. 2D). For the long major repeat (^1^NANPNANPNANPNANPNANPNANP^24^), residues were numbered from 1 to 24, although no electron density was visible for Asn^17^ to Pro^24^ and was not well defined for Asn^1^ (Fig. 2E). The experimental electron density for all of these peptides bound to MAM01 Fab is shown in Fig. S1A. MAM01 bound to all CSP-derived peptides in a similar manner, and their superimposition revealed that all of the peptides adopted secondary structural motifs corresponding to type I β-turn conformations with high similarity in the central type I β-turn (Fig. 2F, Fig. S1B and C). The core of the major repeat region displayed an identical conformation to the core of the minor repeat region, with all epitopes forming similar hydrogen bond interactions with heavy chain Arg^52^, Tyr^53^, Gln^96^, Gln^100D^, and Tyr^100E^, and light chain Gly^30^, Tyr^31^, Asp^32^, and Trp^96^ in the MAM01 paratope (Fig. 2G). However, fewer hydrogen bonding interactions were observed for MAM01 Fab complexed with the junctional region peptide in CDRL1 and CDRH3 (Fig. 2H).

Furthermore, we compared the crystal structure of Fab MAM01 with short major repeat region peptide (NPNA_3_) with our previously determined antibody Fab 224 complex with NPNA_3_ (PDB ID:6WFY) (28) to investigate whether any structural conformational changes were observed after the antibody modifications (Fig. 3A and B). MAM01 Fab with short major repeat region peptide (NPNA_3_) crystallized in a P2_1_2_1_2_1_ space group, whereas Fab 224 in complex with NPNA_3_ crystallized in a P2_1_ space group, each with a single complex in the crystal asymmetric unit (asu). Alignment of Fab MAM01 and Fab 224 complexes with NPNA_3_ yields a low RMSD of 0.15 Å for all atoms in the Fab variable domains (Fig. 3C).

**Figure 3.**
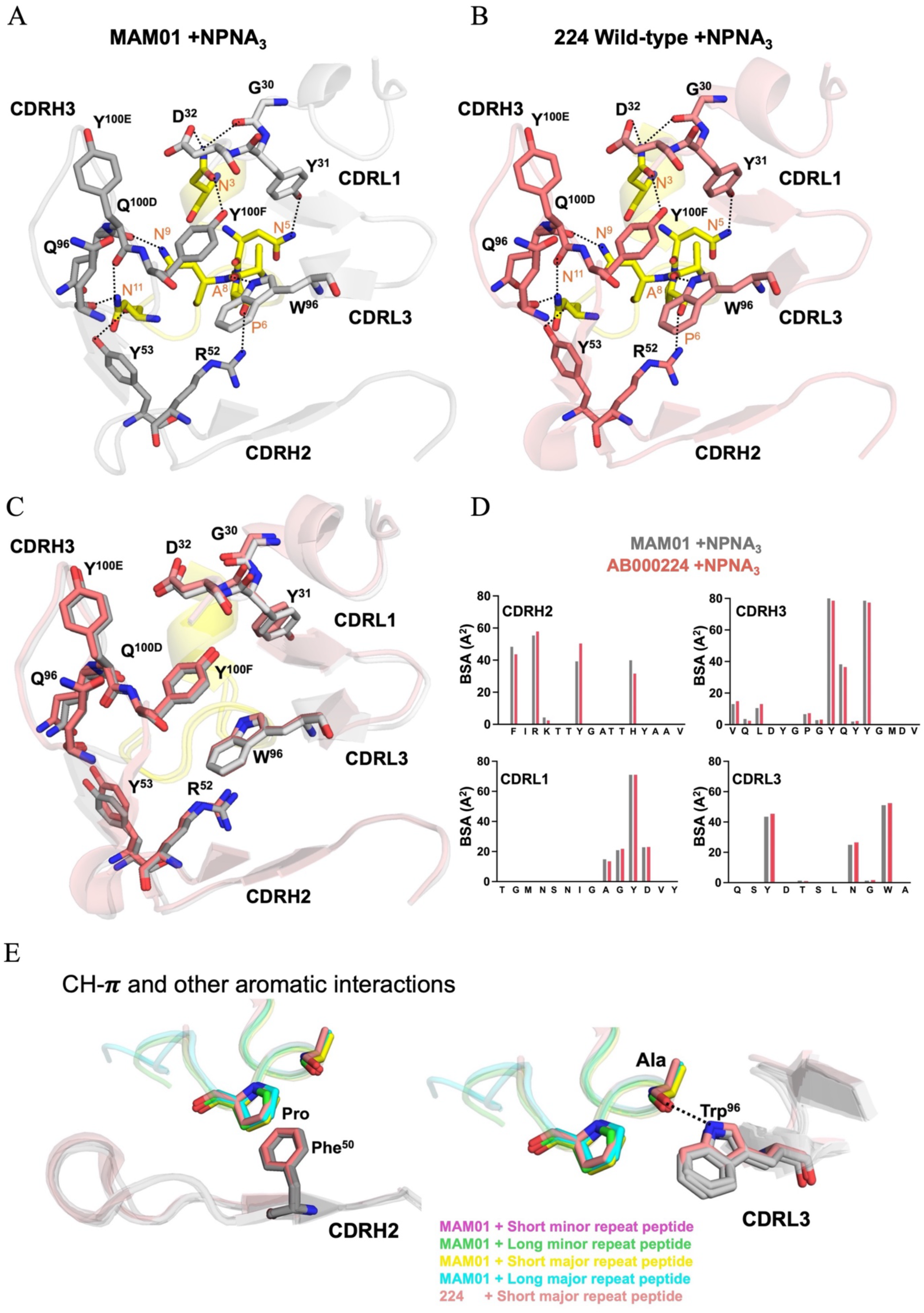
Conserved interactions of AB000224 (224) and MAM01 Fabs. A) Crystal structure of MAM01 with NPNA_3._ B) Crystal structure of Fab 224 with NPNA_3_ (PDB ID: 6WFY). Only CDRs involved in peptide interactions are shown. Residues forming hydrogen bonds are shown as sticks. C) Structural alignment of Fabs 224 (salmon) and MAM01 (grey) with NPNA_3_ (yellow). D) Bar plot showing buried surface area contributions of CDRs H2, H3, L1, and L3 in Fab 224 (salmon) and MAM01 (grey). E) Superimposed view of aromatic residues in Fabs 224 (PDB ID: 6WFY) and MAM01 interacting with CSP-derived peptides in different regions.

The short major repeat peptide NPNA_3_ binds to Fab 224 and MAM01 in a similar type I β-turn conformation (Fig. S1C). The interactions with CDR loops and hydrogen-bond patterns are conserved in both 224 and engineered MAM01 Fabs. The buried surface area (BSA) for the Fab MAM01-NPNA_3_ complex is primarily contributed by the heavy chain (446 Å^2^ on the Fab and 496 Å^2^ on the peptide), similar to Fab 224-NPNA_3_ complex (443Å^2^ on the Fab and 496 Å^2^ on the peptide). Light-chain contributions of MAM01 are mediated by CDRL1 and CDRL3 (BSA: 252 Å^2^ on the Fab and 264 Å^2^ on the peptide), comparable to those in the Fab 224-NPNA_3_ complex (BSA: 256 Å^2^ on the Fab and 263 Å^2^ on the peptide) (Fig. 3D). The highest contributions to the BSA come from P^10^, N^11^, N^3^, N^7^ in ^3^NANPNANPNA^12^ in both crystal structures (Fig. S2).

Furthermore, the MAM01-peptide complexes have similar interactions of Trp-Pro and Trp-hydrogen bonds as in Fab 224 with the NPNA_3_ peptide. Fab 224 and MAM01 both engage in CH/π interactions with germline-encoded Phe^50^ in CDRH2 and similar hydrogen bond interactions between conserved Trp^96^ in CDRL3 and the alanine backbone (Fig. 3E).

### Crystal structures of mAb 7088 Fab and MS-1805 Fab with junctional and short major repeat regions

Although mAb AB007088, derived from the IGHV3-33/IGKV1-5 germline gene, was engineered to MS-1805 as an alternative candidate for malaria prevention, structural data for 7088 Fab, both alone and in complex with CSP-derived peptides, were previously unavailable. Here, we determined crystal structures of Fab 7088, and its engineered counterpart MS-1805, with peptides derived from the short major repeat (NPNA_3_) and N-terminal junctional region. Crystals of Fab 7088 diffracted to 3.20 Å and 2.0 Å resolution for the N-terminal junctional and short major repeat regions, respectively (Fig. 4A, Table S3), and crystals of Fab MS-1805 Fab diffracted to 1.74 Å for the N-terminal junctional region and 2.25 Å for the short major repeat region (Fig. 4B, Table S3).

**Figure 4.**
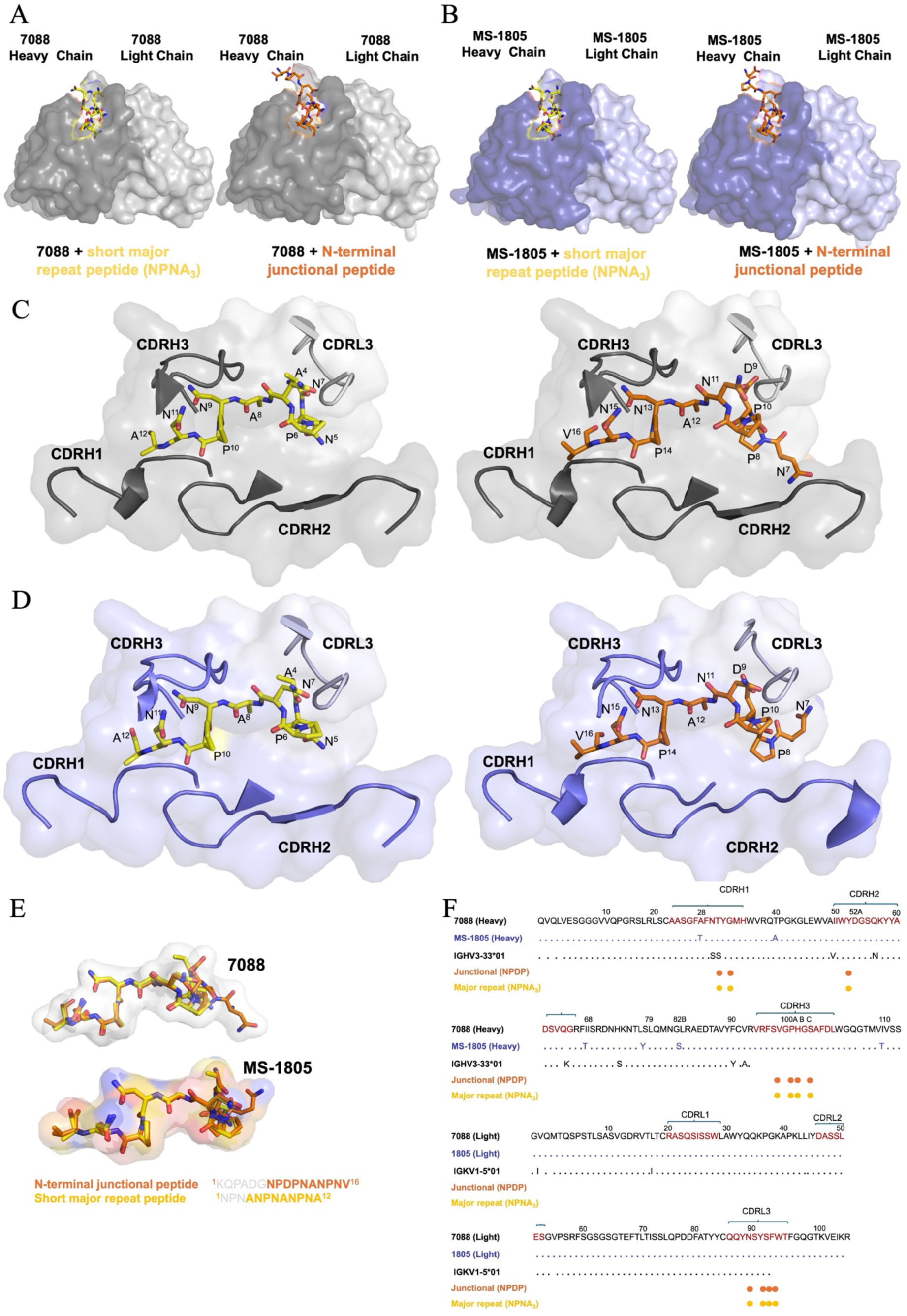
Crystal structures of Fab AB007088 (7088) and MS-1805 with CSP-derived peptides. A) Surface representation of Fab 7088 (heavy chain: grey, light chain: white) with short major repeat (yellow sticks) and junctional (orange sticks) peptides. B) Structure of engineered Fab MS-1805 (heavy chain: dark blue, light chain: light blue) with the same peptides. C) CDRs involved in interactions with peptides in Fab 7088 (grey) and MS-1805 (blue), are shown as cartoons. E) Superimposition of N-terminal junctional and major repeat peptides of 7088 and MS-1805 embedded in transparent surface rendering are represented as sticks and colored with respect to the sequences observed in electron density. F) Sequence of Fabs 7088 (black) and MS-1805 (blue) with CDR residues interacting with each peptide marked as closed circles, color-coded by peptide region.

In Fab 7088 and Fab MS-1805 with NPNA_3_, electron density was not observed for the initial three residues (Asn^1^ to Asn^3^). Similarly, for Fab 7088 and MS-1805 with the N-terminal junctional peptide (^1^KQPADGNPDPNANPNV^16^), there was no electron density for the initial six residues (Lys^1^ to Gly^6^) (Fig. 4C and D and Fig. S3A), suggesting that the KQPADG region is disordered and not essential for binding. The Fab 7088 with short major repeat region (NPNA_3_) crystallized in a P4_3_2_1_2 space group, while Fab 7088 with the N-terminal junctional region crystallized in space group P2_1_2_1_2_1_ with a single complex in the crystal asymmetric unit (asu). In contrast, engineered Fab MS-1805 with NPNA_3_ and the N-terminal junctional region crystallized in a P2_1_ space group with a single complex in the crystal asymmetric unit (asu). Interactions between Fab 7088 and engineered Fab MS-1805 with their CSP-derived peptides are mainly mediated through heavy chain CDRH1, H2, and H3, with additional interactions via CDRL3. Both Fab 7088 and its engineered counterpart Fab MS-1805 bound their peptides in a similar Asn pseudo 3_10_-turn and type I β-turn conformation (Fig. 4E and Fig. S3B).

Fab 7088 and MS-1805 complexes with the major repeat region peptide showed an almost identical conformation to those complexed with the N-terminal junctional region, with all epitopes forming similar hydrogen bond interactions. Key conserved interactions are made with heavy chain Thr^31^, Gly^33^, Tyr^52A^, Ser^98^, Gly^100^, Gly^103^, Pro^101^, and light chain Asn^92^, Tyr^94^, Ser^95^ and Phe^96^ in the paratope of Fab 7088 and MS-1805 (Fig. 4F).

We further superimposed the crystal structure of Fab MS-1805 with Fab 7088 in complex with both short major repeat (NPNA_3_) and N-terminal junctional region peptides to investigate potential differences in epitope interactions or conformation. The epitope interactions with CDR loops were similar, with conserved hydrogen-bonding interactions in both 7088 and MS-1805 Fabs with NPNA_3_ (Fig. 5A). The buried surface area (BSA) for Fab MS-1805-NPNA_3_ complex is predominantly contributed by heavy chain (443 Å^2^ on the Fab and 560 Å^2^ on the peptide), like the Fab MS-7088-NPNA_3_ complex (421Å^2^ on the Fab and 526 Å^2^ on the peptide). Contributions from the light chain are mediated exclusively by CDRL3 (189 Å^2^ on the Fab and 248 Å^2^ on the peptide), like those in the Fab MS-7088-NPNA_3_ complex (BSA is 182 Å^2^ on the Fab and 234 Å^2^ on the peptide) (Fig. 5B).

**Figure 5.**
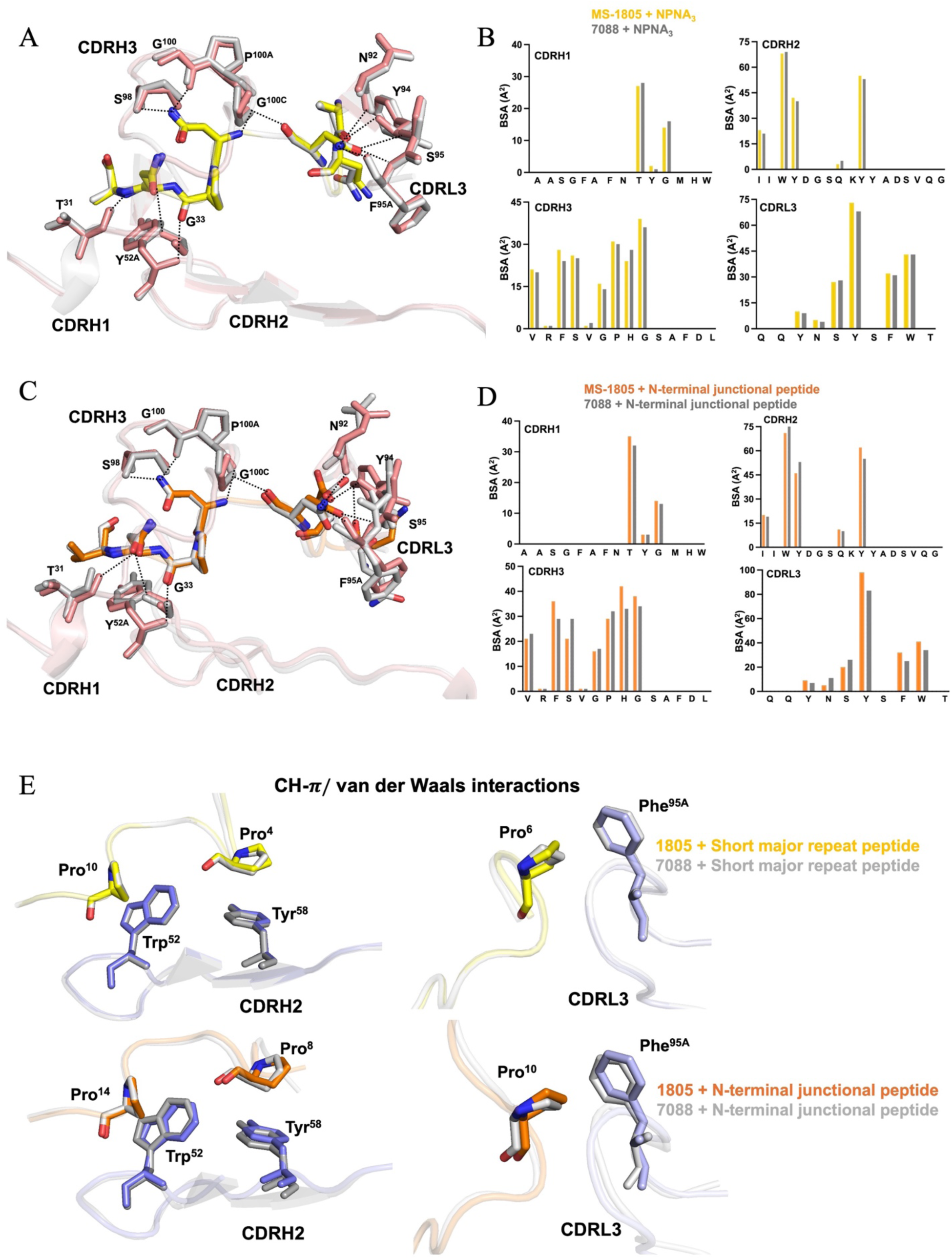
Conserved hydrogen bonding in Fabs 7088 and MS-1805. A) Structural alignment of Fabs 7088 and MS-1805 with NPNA_3_. CDR residues involved in hydrogen bonding are shown as sticks. B) Bar plot of buried surface area contributions of CDRs H1, H2, H3, and L3 in Fab 7088 (grey) and MS-1805 (yellow). C) Structural alignment of Fab 7088 and MS-1805 with the N-terminal junctional peptide (orange). D) Bar plot showing buried surface area contributions of CDRs H1, H2, H3, and L3 for Fabs 7088 (grey) and MS-1805 (orange). E) Superimposed aromatic residues in Fabs 7088 (grey) and MS-1805 (blue) with CSP-derived peptides.

Notably, we observed slight differences in the initial residues ^7^NPD^9^ of the N-terminal junctional epitope in the MS-1805 complex, which interacts with CDRH3, compared to Fab 7088 complexed with the same peptide. This minor difference may be due to the lower resolution of Fab 7088 structure, and less resolved density of the peptide and CDRH3 loop. However, the epitopes interactions and hydrogen bonds with CDRs were conserved in both 7088 and 1805 Fabs complexes with the N-terminal junctional peptide (Fig. 5C). The BSA for the Fab MS-1805 with N-terminal junctional peptide complex is primarily contributed by heavy chain (463 Å^2^ on the Fab and 587 Å^2^ on the peptide), similar to that for the Fab 7088 complexed (446 Å^2^ on the Fab and 582 Å^2^ on the peptide). The light chain BSA is mediated solely by CDRL3 (208 Å^2^ on the Fab and 251 Å^2^ on the peptide), similar to Fab 7088 (198 Å^2^ on the Fab and 240 Å^2^ on the peptide) (Fig. 5D).

Most of the BSA contributions from heavy chain come from residues ^10^PNANPNV^15^ in both Fabs 7088 and MS-1805 with the N-terminal junctional peptide (Fig. S4A). Similarly, in Fabs 7088 and MS-1805 with NPNA_3_, the majority of the BSA from the heavy chain is from ^6^PNANPNA^12^ (Fig. S4B), indicating that the minimal epitope PNANPN is necessary for both junctional and major repeat peptides.

Furthermore, the MS-1805 peptide complex exhibited similar Trp-Pro and other aromatic interactions as observed in Fab 7088 with NPNA_3_ and junctional region peptides, with both complexes engaging in CH/π interactions with germline-encoded Trp^52^ and Tyr^58^ in CDRH2 (Fig. 5E). Interestingly, we noted that Phe^95A^ in both 7088 and engineered MS-1805 Fab participates in CH/π interactions with proline in the short major repeat and N-terminal junctional epitopes. This interaction is distinct from the other known Fabs that are encoded by the IGHV3-33/IGKV1-5 germline genes (27, 29).

### Highly protective mAb Fab 7088 and engineered variant Fab MS-1805 features an extended CDRL3 compared to other antibodies encoded by the IGHV3-33/1GKV1-5 germlines

Previous studies have shown that antibodies capable of cross-reacting with the NANP repeat peptide are often encoded by IGHV3-33 paired with IGKV1-5 (27). Several crystal structures of Fabs derived from the IGHV3-33/ IGKV1-5 germline have been determined. To better understand the molecular recognition of antibodies from these specific germlines, we compared crystal structures of MS-1805 with NPNA_3_ to other antibodies, including Fabs 850, 1210, 2541, and 2243, which were also in complex with major repeat region peptides (29, 30) (Fig. 6).

**Figure 6.**
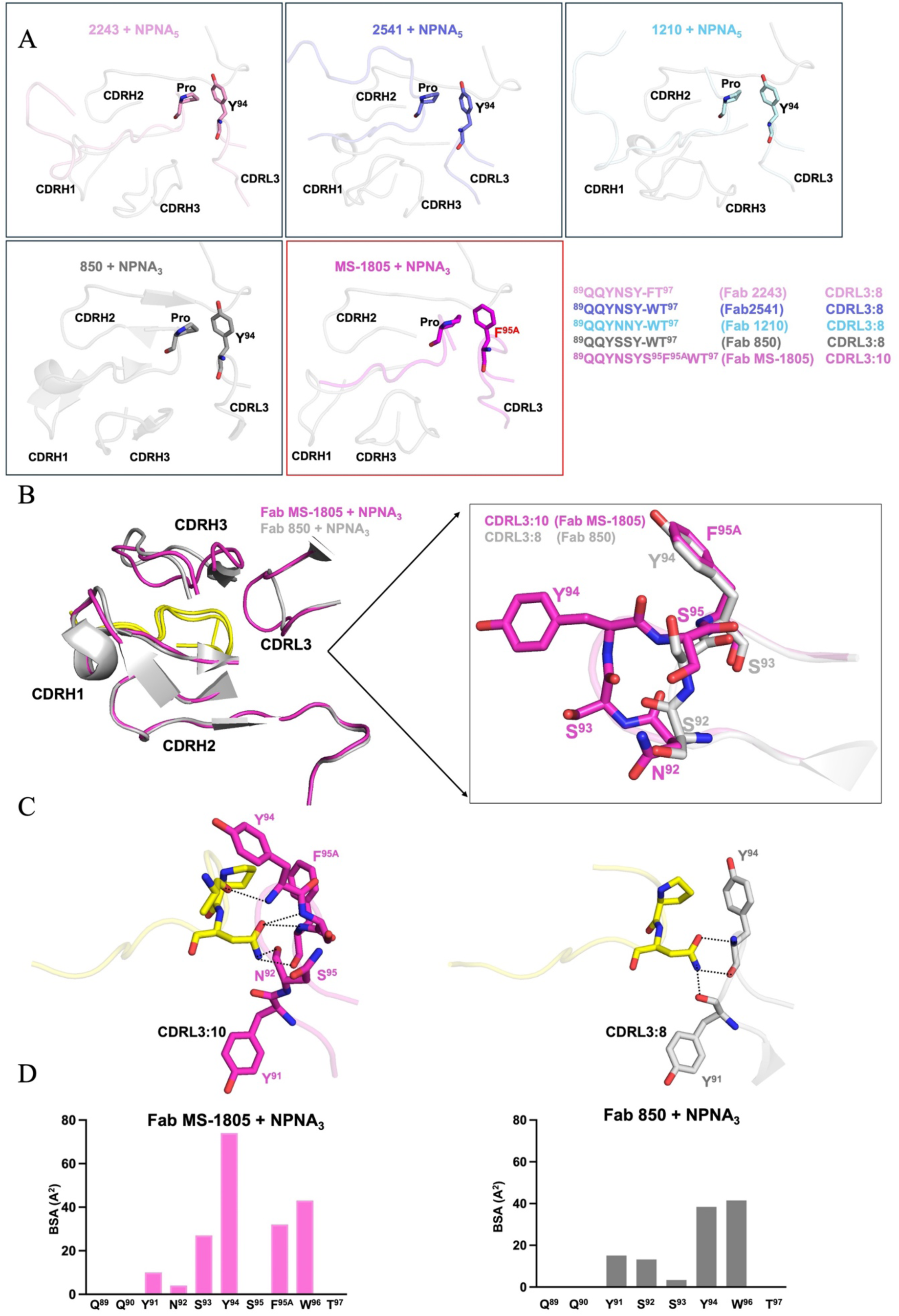
Structural comparison of Fabs to CSP encoded by IGHV3-33/IGKV1-5 germline genes. **A)** Semi-transparent cartoon representation of Fab MS-1805 (magenta) with other IGHV3-33/IGKV1-5 Fabs 850 (grey), 1210 (cyan), 2541 (slate blue), and 2243 (pink), each complexed with peptides (yellow). CH/π interactions of Phe^95A^ in CDRL3 of Fab MS-1805 and Tyr^94^ in CDRL3 of Fab 850 (PDB ID: 7UYM), Fab 1210 (PDB ID: 6D01), 2541 (PDB ID: 6ULE), and 2243 (PDB ID: 6O23) engaged with proline are shown as sticks and highlighted in their respective colors. **B)** Superposition of Fab MS-1805 and Fab 850 with NPNA_3_, highlighting their CDRL3 conformations. **C)** Hydrogen-bond interactions (black dashes) between Fab MS-1805 (magenta) and Fab 850 (grey) with the short major repeat peptide (yellow). **D)** Bar plot shows the individual residues of CDRL3 that contribute to the buried surface area of Fab MS-1805 (colored in magenta) and Fab 850 (colored in grey). Crystal structure of Fab 850 complexed with NPNA_3_ was obtained from a previous study (PDB ID: 7UYM)

All of these Fabs exhibit similar conformations in their CDRH1, CDRH2 and CDRH3 regions, suggesting conserved interactions with their respective epitopes. However, a key difference was that MS-1805 possesses a CDRL3 of 10 amino acids (CDRL3:10), whereas CDRL3 of other related antibodies in this germline is typically 8 amino acids (CDRL3:8). Interestingly, while a tyrosine at position 94 in CDRL3 of the other Fabs forms CH/π interactions with proline, MS-1805 has a phenylalanine at position 95A in CDRL3 that engages in similar CH/π interactions (Fig. 6A). To further understand how the extended CDRL3 in accommodated in MS-1805, we superimposed its crystal structure with that of Fab 850 with NPNA_3_ (PDB ID: 7UYM) (30). In Fab MS-1805, CDRL3 is extended at its tip by Ser and Tyr at positions 93 and 94, and Ser^95^ and Phe^95A^ now occupy positions corresponding to Asn^93^ and Tyr^94^ in Fab 850 (Fig. 6B). Additionally, CDRL3:8 in Fab 850 has fewer hydrogen-bond interactions with its epitope compared to CDRL3:10 in Fab MS-1805, where Ser^95^ and Phe^95A^ form additional interactions (Fig. 6C). Furthermore, the BSA contributed by CDRL3 in MS-1805 is 189 Å^2^ on Fab and 248 Å^2^ on peptide, substantially higher than the BSA contributed by CDRL3 in Fab 850 (111 Å^2^ on Fab and 165 Å^2^ on the peptide) (Fig. 6D). Overall, the crystal structure of MS-1805 with a longer CDRL3 showed that this extended loop plays a key role in the unique binding profile and specificity of this antibody. These findings suggested that the extended length of CDRL3 in MS-1805 may contribute to its superior affinity and specificity by facilitating optimal interactions with the epitope, potentially offering advantages over other IGHV3-33/IGKV1-5 germline encoded antibodies in therapeutic applications.

### The unliganded structure of Fab MAM01 and MS-1805 show small conformational changes in their CDR side chains upon peptide binding

We determined unliganded crystal structures of MAM01 and MS-1805 at resolutions of 2.08 Å and 1.77 Å, respectively, to investigate any conformational changes upon binding to CSP peptides (Table S4). We superimposed the unliganded structure of MAM01 with its peptide-bound forms. We observed changes in the orientation of some side chains in the CDRs in Fab MAM01 unliganded structure compared to the liganded structures. The aromatic side chains of Tyr^53^ in CDRH2, Tyr^100E^ and Tyr^100F^ in CDRH3, and Trp^96^ in CDRL3 of MAM01 showed some movement, mainly in their rotamers, when bound to the peptides (Fig. 7A). Similar side-chain reorientations were seen in unliganded Fab 224 when compared to its NANP_3_ bound structure, with notable movements in Tyr^100E^, Tyr^100F^ in CDRH3, Trp^96^ in CDRL3, and Arg^52^ in CDRH2 (Fig. S5). In the unliganded MS-1805 structure, slight shifts were observed for tip of CDRH3 residues 98-100C that include peptide contact residues Ser^98^, Pro^101^ and Gly^100C^ (Fig. 7B).

**Figure 7.**
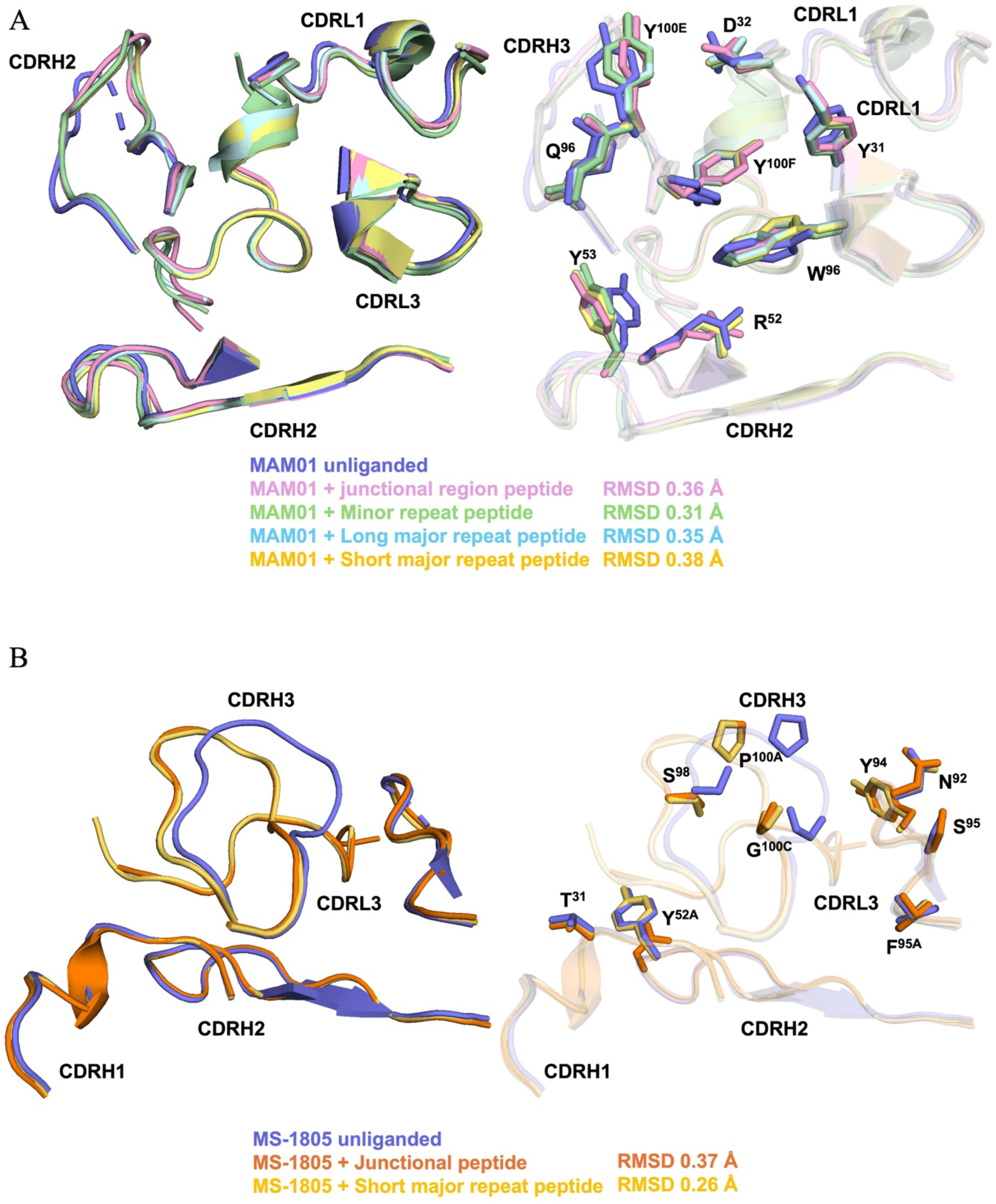
Unliganded structures of engineered Fabs MAM01 and MS-1805. **A)** Structural alignment of unliganded MAM01 (blue) with its peptide-bound forms: junctional (pink), minor (green), short major repeat (yellow), and long major repeat (cyan). **B)** Alignment of unliganded MS-1805 (blue) with peptide-bound forms: junctional (orange) and short major repeat (yellow). Only side chains of CDR residues involved in hydrogen bonds are shown as sticks. Root mean square deviation (RMSD) values are shown (Å) for each alignment.

## Discussion

This study focused on the structural analysis of highly protective and cost-effective engineered monoclonal antibodies MAM01 and MS-1805, which are derived from different heavy and light chain germline genes and bind to different regions of CSP-derived peptides.

MAM01 has been and is currently in clinical trials (US, Uganda) and is being evaluated for use as a seasonal prophylactic treatment for malaria, particularly in pediatric populations in malaria-endemic regions. From crystal structures and binding experiments, we observed that MAM01 Fab binds to several epitopes, including the junctional region, minor repeat region, and major repeat region with similar secondary type I β-turn structural conformations. Comparing the structures of the engineered Fab MAM01 with Fab 224 complexed with an NPNA_3_ repeat revealed that specific mutations in framework region of the MAM01 variable domains do not cause any conformational changes or disrupt interactions with its epitope. Other highly protective engineered mAbs L9LS and CIS43LS, which bind specifically to minor (NVDP) and junctional (NPDP) regions, respectively, are also in clinical trials (18). It will be interesting to investigate and compare the in vivo study of the engineered antibodies MAM01, L9LS and CIS43LS, which would provide valuable insights into their efficacy against malaria.

Another monoclonal antibody AB007088 is derived from a predominant lineage IGHV3-33/IGKV1-5 elicited by RTS,S vaccinees and showed a superior protection against malaria in a parasitemia challenge model (22). This mAb was engineered, renamed as MS-1805, and initially selected as an alternative candidate to MAM01, but no clinical trials have been initiated. Crystal structures of Fab 7088 and engineered MS-1805 in complex with N-terminal junctional peptide and short major repeat (NPNA_3_) reveal almost identical conformations and similar hydrogen bonding interactions with their epitopes. The majority of crystal structures to the NPNA repeat region are encoded by the germline lineage IGHV3-33 (28). Fab MS-1805 has similar germline-encoded Trp^52^ and Tyr^58^ CH/π interactions with proline. Interestingly, we observed that CDRL3 of Fab MS-1805 is longer than the other known antibodies to CSP NPNA repeats encoded by the same light chain lineage. The longer CDRL3 of MS-1805 showed an increase in the buried surface area and interactions with its epitopes, consistent with its higher affinity compared to other antibodies encoded by the same germline genes. It will be interesting to determine how the longer CDRL3 might affect its functional properties in a biological or therapeutic setting. Furthermore, crystal structures of IGHV3-33/IGKV1-5 Fab 1210, Fab 2243, Fab 2541, and Fab 850 in complex with NANP_5_ showed affinity-matured homotypic interactions (26, 29, 30). However, in our crystal structure, MAM01 in complex with the long major repeat (NANP_6_) does not form homotypic interactions. Whether Fab 7088 and its engineered variant MS-1805 are capable of such interactions remains to be determined. Although crystallization trials of 7088 and MS-1805 with NANP_6_ were unsuccessful, further structural studies will be required to establish whether this highly protective antibody can engage in homotypic contacts (31).

Although both AB000224 (MAM01) and AB007088 (MS-1805) monoclonal antibodies were isolated from B cells of protected RTS,S vaccinees, they recognize epitopes in addition to NPNA repeats, including junctional and minor repeat regions, which are not present in RTS,S. From our structural analysis, we observed that the majority of BSA contribution is mediated by the PNANPN sequence that is present in the junctional, minor and major repeat regions. Although these antibodies were likely affinity-matured against the major repeat, they nevertheless exhibit promiscuous binding to PNANPN in the context of junctional and minor repeat regions. Structural analysis of MAM01 and MS-1805 showed that these cross-reactive human mAbs recognize distinct epitopes of *P. falciparum* circumsporozoite protein, each epitope adopting a specific secondary structural motif, corresponding either to type I β-turn or Asn pseudo 3_10_ turn conformations.

The NPDP junctional and NVDP minor repeat motif of PfCSP, which is targeted by both protective mAbs, represents a neutralizing epitope that is not present in the RTS,S vaccine. These findings confirm that incorporating these epitopes could enhance the effectiveness of future malaria vaccines (14, 15, 29, 32, 33). Overall, these engineered antibodies, MAM01 and MS-1805, are a promising advance in malaria prevention due to their mechanism of action, successful manufacture, and long-acting formulation.

## Materials and Methods

### Protein expression and purification

Antibody sequences of heavy chain and light chain variable regions were obtained from Atreca and are available in (22). The Fab genes of MAM01, 7088, and MS-1805 were codon optimized for mammalian expression and cloned into the expression vector PHCMV3 by GenScript. The Fabs were expressed in ExpiCHO cells and purified using a HiTrap Protein G HP column (GE Healthcare) followed by size exclusion chromatography (Superdex 200 16/90; GE Healthcare) in 1X Tris-buffered saline buffer (TBS : 50 mM Tris pH 7.8, 140 mM NaCl).

### Bio-layer interferometry

Binding kinetics of Fabs 224, MAM01, 7088, and MS-1805 to different epitopes of PfCSP were determined by Biolayer Interferometry (BLI, Octet Red; Pall ForteBio). All biotinylated peptides were ordered from Innopep Inc. and diluted to 10 μg/mL in kinetics buffer (1X PBS containing 0.01% BSA and 0.002% Tween 20 and loaded on streptavidin biosensors. Loading of biotinylated peptides was performed for 300s, followed by 40s baseline in kinetics buffer. Association with Fabs (serial dilution from 200 nM, 100 nM, 50 nM, 25 nM, to 12.5 nM) was performed for 300s, followed by a dissociation step in buffer for 300s. Background subtraction was performed in buffer alone to assess non-specific binding. Kinetics of BLI data were measured using Octet data analysis software, version 12.2, and fitted with a 1:1 binding model.

### Fab-peptide complexes

All the CSP derived peptides had protection of their amine and carboxyl groups at the N-term and C-term with acetyl and amine groups: Ac-KQPADGNPDPNANPNV-NH_2_ (N-terminal junctional region peptide), Ac-NPDPNANPNVDPNANP-NH_2_ (Junctional region peptide), Ac-NVDPNANPNVDPNANPNVDP-NH_2_ (Minor repeat region peptide), Ac-NPNANPNANPNA-NH_2_ (Short major repeat region peptide) and Ac-NANPNANPNANPNANPNANPNANP-NH_2_ (Long major repeat region peptide). Peptides were ordered from Innopep Inc. with high (98%) purity. MAM01Fab was concentrated to 11 mg/ml in 1X TBS using 10K Amicon® ultra centrifugal units and further mixed with junctional, minor, and major repeat peptides in a 1:5 molar ratio of Fab to peptide. Similarly, 7088 Fab and MS-1805 Fab were concentrated to 10 mg/ml in 1X TBS and mixed with N-terminal junctional and short major repeat peptides in a 1:5 molar ratio of Fab to peptide and stored at 4 °C overnight.

### X-ray crystallization

All crystal screening was performed using our high-throughput robotic CrystalMation system (Rigaku) at TSRI. Crystals were grown using the sitting drop vapor diffusion method with a reservoir solution of 35 µL with each drop consisting of 0.1 µL of protein and 0.1 µL precipitant. Crystals for Fab MAM01 complexed with the junctional peptide were grown in 0.085 M Tris pH 8.5, 0.17 M sodium acetate, 15% (v/v) glycerol, and 25.5 % (w/v) polyethylene glycol 4000. Fab MAM01 with the minor repeat peptide were grown in 0.2 M potassium acetate, pH 7.8 and 20% (w/v) polyethylene glycol 3350 and cryoprotected in 10% ethylene glycol. Fab MAM01-NPNA_3_ cocrystals were grown in 0.1 M Bicine pH 9.0 and 65% (v/v) 2-methyl-2,4-pentanediol. Crystals of the Fab MAM01-NANP_6_ complex were grown in 0.04 M potassium dihydrogen phosphate, 20% (v/v) glycerol and 16% (w/v) polyethylene glycol 8000. Crystals for MAM01-apo were grown in 0.1 M MES pH 6.0 and 30% (w/v) polyethylene glycol 6000 and cryoprotected in 10% PEG 200. Crystals for Fab 224-apo were grown in 0.17 M ammonium sulfate, 15% (v/v) glycerol, 25.5% (w/v) PEG4000. All crystals for Fab MAM01 and Fab 224 complexed with or without their peptides appeared within 7 days at 293 K.

Crystals for 7088 Fab with junctional region peptide were grown in 0.1 M Tris pH 8.0, 1M lithium chloride and 10% (w/v) polyethylene glycol 6000 and cryoprotected in 15% ethylene glycol while for 7088-NPNA_3_ complex were grown in 0.2 M sodium chloride and 20% (w/v) polyethylene glycol 3350 and cryoprotected in 15% ethylene glycol. Co-crystals of MS-1805 Fab with junctional peptide were grown in 0.1 M sodium citrate, 20% (v/v) 2-propanol, and 20% (w/v) polyethylene glycol 4000 and Fab 1805 with NPNA_3_ were grown in 0.2 M zinc acetate and 20% (w/v) polyethylene glycol 3350; both sets of crystals were cryoprotected in 10% ethylene glycol. Crystals for Fab MS-1805 apo were grown in 0.095 M sodium citrate pH 5.6, 19% (v/v) 2-propanol, 5% glycerol and 19% (w/v) polyethylene glycol 4000. All crystals for Fab 7088 and 1805 with or without peptides appeared within 14-21 days at 293 K.

### X-ray data collection and structure determination

X-ray diffraction data for MAM01 Fabs with or without peptides were collected at Stanford Synchrotron Radiation Lightsource (SSRL) beamline BL12-1 (Tables S2 and S4). For Fabs 7088 and MS-1805 with or without peptides, x-ray diffraction data were collected at National Synchrotron Light Source II beamline 17-ID-1 (highly automated macromolecular crystallography - NSLS-II AMX) (Tables S3 and S4). X-ray diffraction data for Fab 224 apo were collected at National Synchrotron Light Source II beamline 17-ID-2 (frontier micro focusing macromolecular crystallography - NSLS-II FMX). All diffraction data were processed and scaled using the HKL-2000 package (34). The structures of MAM01 Fab were solved by molecular replacement with PHASER (35) using Fab 224 (PDB ID: 6WFY) as a search model. The structures of 7088 Fab were solved by molecular replacement with PHASER using a homology model generated with SWISS-MODEL (36, 37), and subsequent MS-1805 Fab structures were solved using Fab 7088 as the search model. All Fab structures were refined using phenix.refine combining with model building cycles in Coot (38, 39) The peptides were manually fitted into the Fo-Fc electron density maps with multiple rounds of refinement in phenix.refine (38). The residue numbering of the Fabs follows Kabat nomenclature and structure validation was done with MolProbity (40). Buried surface areas were calculated using the program MS (41) and hydrogen bonds were assessed using the program HBPLUS for structure analysis (42).

## Data availability

The authors declare that all the data supporting the findings of this study are available within the manuscript and supplementary information files. The structure factors and coordinates of MAM01 bound to junctional, minor, short major repeat, and long major repeat regions have been deposited in the Protein Data Bank (PDB) with accession codes 9NL1, 9NZF, 9NL0, and 9NKZ, respectively. Similarly, the structure factors and coordinates of 7088 bound to the N-terminal junctional and short minor repeat region and MS-1805 bound to the N-terminal junctional and short minor repeat region have been deposited with PDB accession codes 9DSU, 9DSS, 9DST, 9DSR, respectively. The structure factors and coordinates of 224, MAM01and MS-1805 in apo form have been deposited with PDB accession codes 9NOB, 9NKYand 9NOA. respectively.

## Acknowledgements

We thank Atreca, Inc. for providing the antibody sequences of heavy chain and light chain variable regions of the engineered Fabs MAM01 and MS-1805, as also reported in ref. 22. This work was supported and funded by Gates Foundation grants (INV-004923 and INV-056202) under a collaborative agreement with The Scripps Research Institute. The conclusions and opinions expressed in this work are those of the author(s) alone and shall not be attributed to the Foundation. Under the grant conditions of the Foundation, a Creative Commons Attribution 4.0 License has already been assigned to the Author Accepted Manuscript version that might arise from this submission. Please note that works submitted as a preprint have not undergone a peer review process. We also acknowledge Henry Tien for helping with the automated robotic crystal screening at The Scripps Research Institute, Jacqueline Kirchner of the Gates Foundation for support and helpful comments, and Jared Silverman of the Gates Medical Research Institute for reading the manuscript and providing useful comments.

## Author contributions

Conceptualization: M.J., and I.A.W; Investigation: M.J. and S.A.; Antibody sequences: R.R.K., K.L.W., D.E.E.; Resources: M.J. and G.G.-P; Data curation: M.J. and S.A. Formal analysis: M.J., S.A., R.M. and I.A.W.; Writing-original draft preparation: M.J., and I.A.W.; Writing-reviewing and editing: All authors.

