## Supplemental Figures Tables for "Structural characterization of MAM01 and other cost-effective engineered monoclonal antibodies for malaria prevention"

### Supplementary Figures

**Figure S1. Structural analysis of MAM01 Fab-bound peptides**.

**(A)** 2Fo-Fc electron density maps for junctional (magenta), minor repeat (green), short major (yellow), and long major repeat (cyan) peptides bound to MAM01, contoured at 2.0σ (grey). All peptides are represented as sticks. **(B)** The type I β-turns with connecting hydrogen bonds (black dashes) are shown. **(C)** Ramachandran plots for dihedral angles of MAM01-bound peptides. All peptides have canonical type I β-turns in their NPNA, DPNA, and NPNV motifs based on the Ψ (Psi) angle of residue i+3 in the β-turn (1, 2).

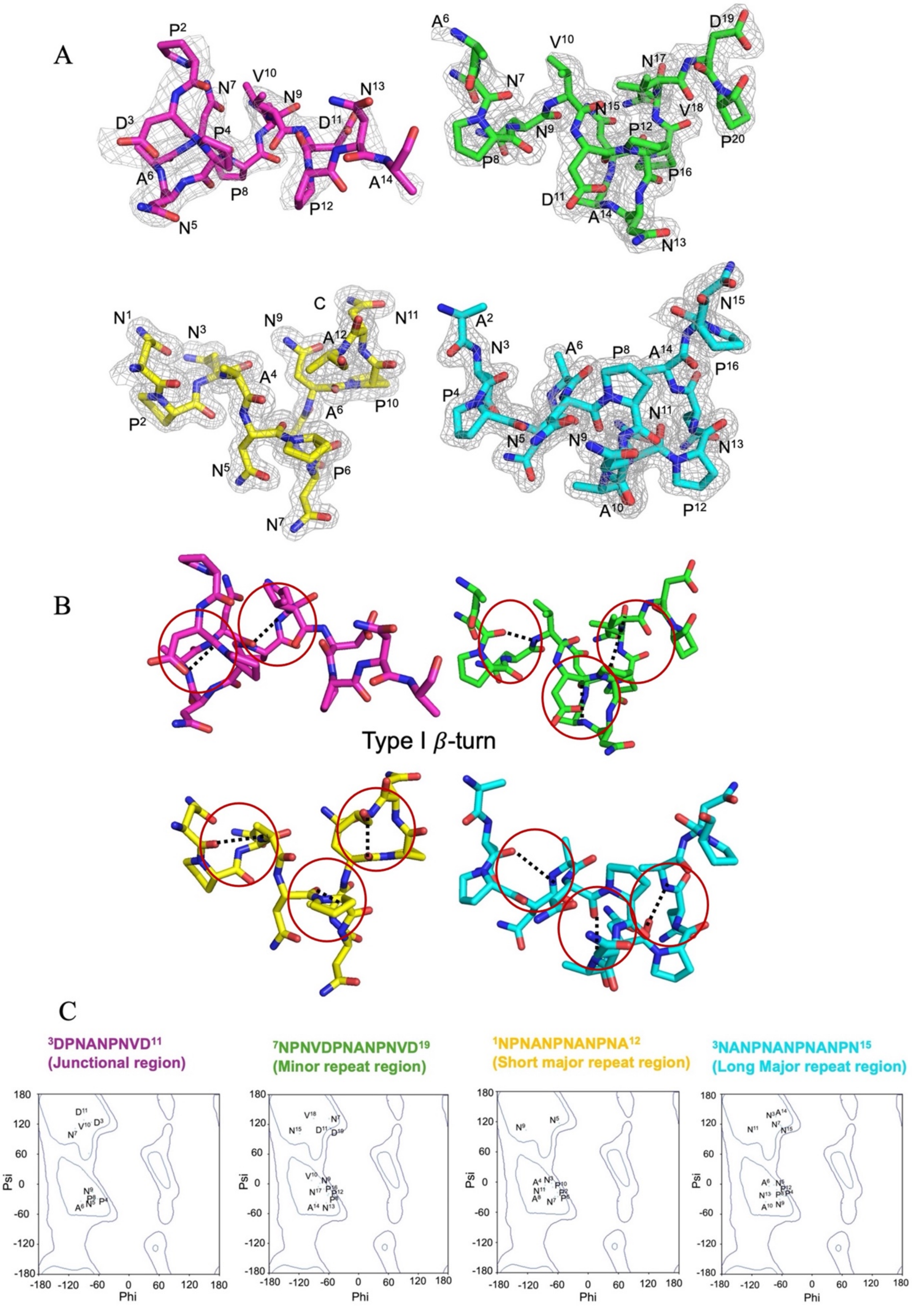

A
B
Type I $\beta$-turn
C
$^{3}$DPNANPNVD$^{11}$
(Junctional region)
$^{7}$NPNVDPNANPNVD$^{19}$
(Minor repeat region)
$^{1}$NPNANPNANPNA$^{12}$
(Short major repeat region)
$^{3}$NANPNANPNANPN$^{15}$
(Long Major repeat region)
Psi
Phi

**Figure S2. BSA for peptides bound to Fab 224 and MAM01.**

Bar plot of buried surface areas of the individual residues for the short major repeat peptide bound to Fabs 224 (colored in grey) and MAM01 (colored in magenta)

A

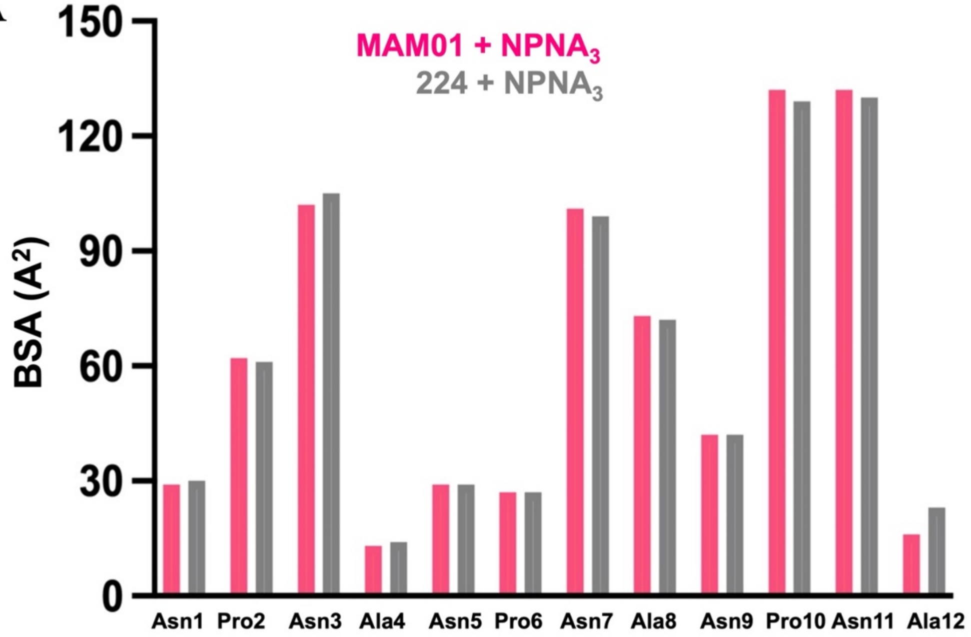

**Figure S3**. **Structural analysis of 7088 and MS-1805 Fab-bound peptides**.

(A) Upper panel shows 2Fo-Fc electron density maps for junctional (orange) and short major repeat peptide (yellow) bound to 7088 Fab, while the lower panel shows 2Fo-Fc electron density maps for peptides bound to MS-1805, contoured at 2.0σ (grey). All peptides are represented as sticks. (B) The type I β-turn (red circle) and Asn pseudo $3_{10}$ turn (blue circle) are shown. The left panel shows the Ramachandran plots for dihedral angles of MS-1805 Fab bound junctional (orange) and short major repeat peptide (yellow) is shown.

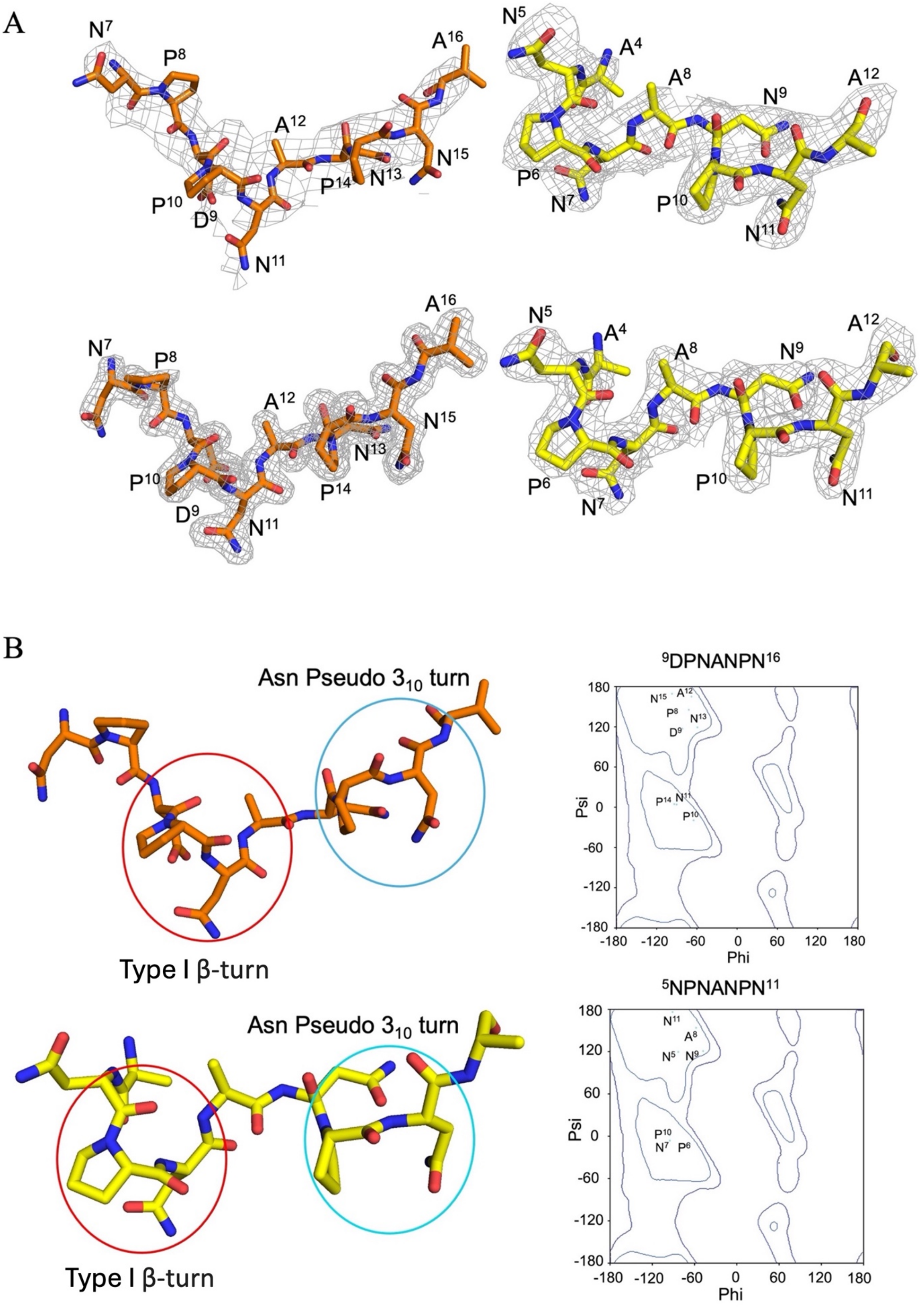
A
N7
P8
D9
P10
N11
A12
N13
P14
N15
A16
A4
N5
P6
N7
A8
N9
P10
N11
A12
B
Asn Pseudo 3₁₀ turn
Type I β-turn
9DPNANPN16
5NPNANPN11
Psi
Phi
180
120
60
0
-60
-120
-180

**Figure S4. BSA for peptides bound to Fabs 7088 and MS-1805.**

A) Bar plot shows buried surface areas of the individual residues for the junctional region peptide bound to Fabs 7088 (colored in grey) and MS-1805 (colored in orange)

B) Bar plot shows buried surface areas of the individual residues for the short major repeat peptide ($NPNA_3$) bound to Fabs 7088 (colored in grey) and MS-1805 (colored in yellow)

A

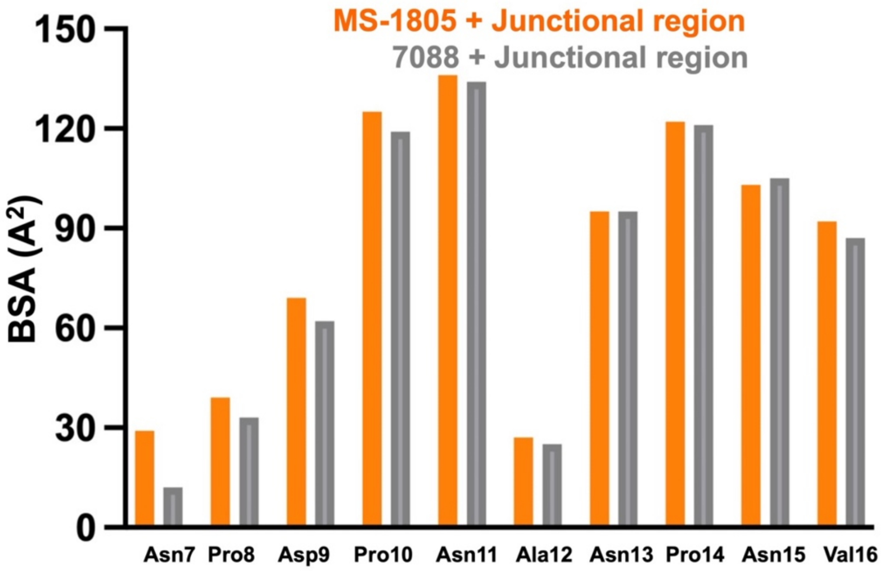


B

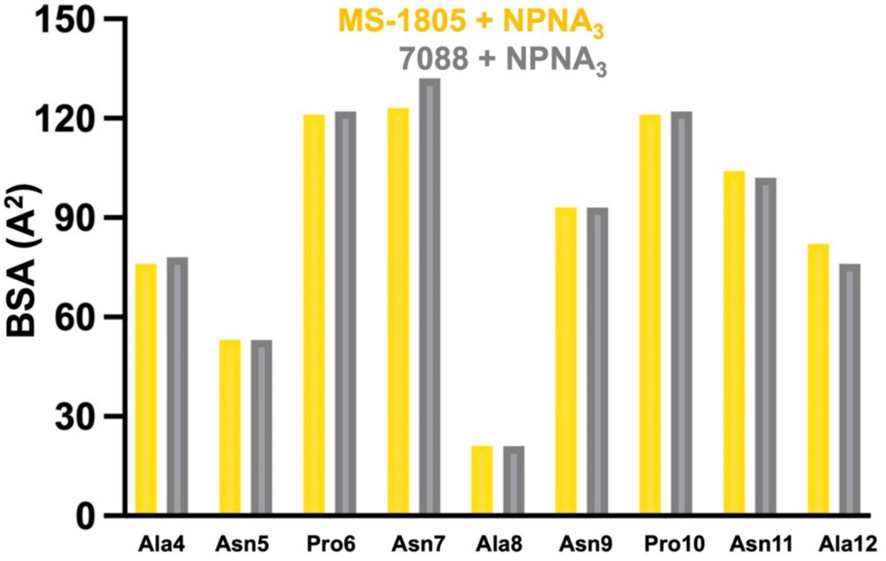

**Figure S5. Crystal structure of unliganded 224 Fab and comparison to its complex with the short major repeat region, related to Figure 7.**

Crystal structure of unliganded 224 Fab (blue) aligned to the 224 Fab in its $NPNA_3$ complex (salmon) are represented as cartoons. Only the side chains of CDRs involved in peptide binding are shown as sticks. Root mean square deviation (RMSD) is indicated (in Angstroms Å) for 194 Cα atoms in the alignment.

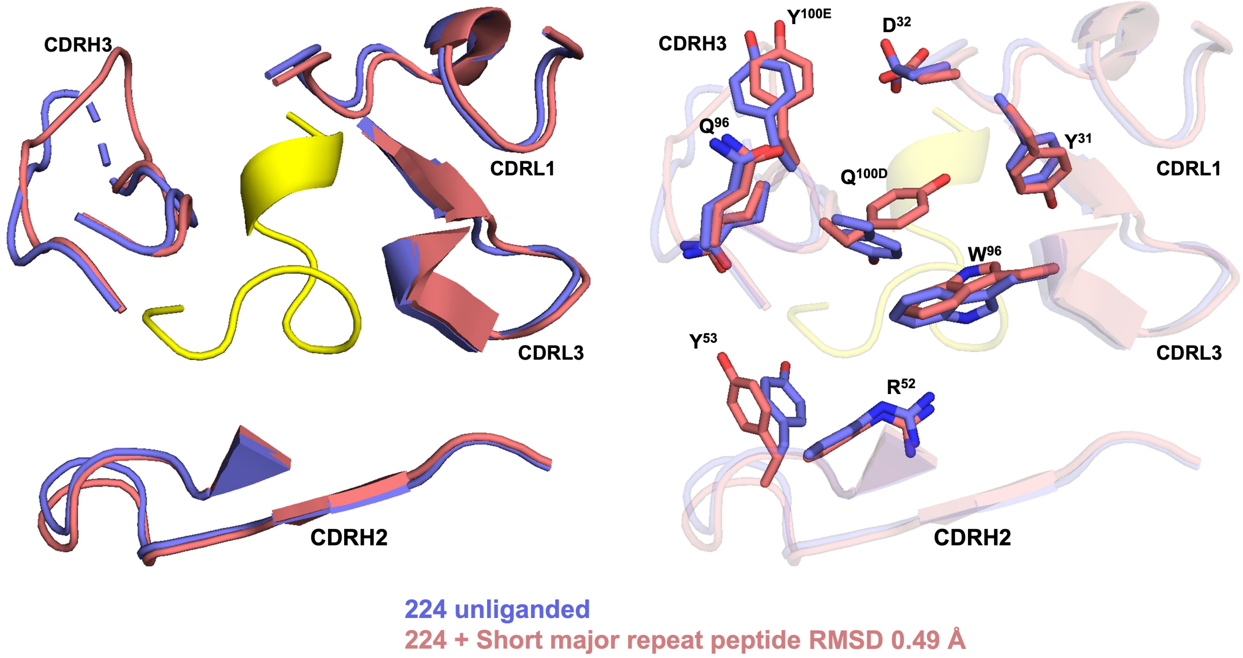

**Table S1. Biolayer interferometry kinetics of Fabs binding to CSP derived peptides**

| Fab | $K_D$ (M) | $K_D$ Error | $k_{on}$ (1/Ms) | $k_{on}$ Error | $k_{off}$ (1/s) | $k_{off}$ Error |
|---|---|---|---|---|---|---|
| **KQPADGNPDPNANPNV (N-term junctional region)** | | | | | | |
| 224 | NB | | | | | |
| MAM01 | NB | | | | | |
| 7088 | 1.55E-08 | 2.83E-11 | 3.13E+04 | 2.84E+01 | 4.86E-04 | 7.69E-07 |
| MS-1805 | 2.39E-08 | 3.35E-11 | 2.96E+04 | 2.69E+01 | 7.09E-04 | 7.55E-07 |
| **NPDPNANPNVDPNANP (Junctional region)** | | | | | | |
| 224 | 7.82E-08 | 1.97E-10 | 6.86E+04 | 1.79E+02 | 4.88E-03 | 4.47E-06 |
| MAM01 | 1.08E-07 | 3.61E-10 | 6.77E+04 | 2.17E+02 | 7.37E-03 | 6.57E-06 |
| 7088 | 1.62E-08 | 4.92E-11 | 3.79E+04 | 6.34E+01 | 6.17E-04 | 1.56E-06 |
| MS-1805 | 2.00E-08 | 5.39E-11 | 3.35E+04 | 5.23E+01 | 6.73E-04 | 1.47E-06 |
| **NVDPNANPNVDPNANPNVDP (Minor repeat region)** | | | | | | |
| 224 | 2.96E-09 | 1.56E-11 | 5.25E+04 | 4.13E+01 | 1.56E-04 | 8.11E-07 |
| MAM01 | 3.53E-09 | 1.46E-11 | 5.08E+04 | 3.69E+01 | 1.79E-04 | 7.31E-07 |
| 7088 | 2.06E-08 | 3.73E-11 | 3.14E+04 | 3.42E+01 | 6.46E-04 | 9.35E-07 |
| MS-1805 | 2.84E-08 | 4.56E-11 | 3.15E+04 | 3.65E+01 | 8.98E-04 | 1.00E-06 |
| **$NPNA_3$ (Major repeat region)** | | | | | | |
| 224 | 3.51E-09 | 1.65E-11 | 6.61E+04 | 6.50E+01 | 2.32E-04 | 1.07E-06 |
| MAM01 | 4.11E-09 | 1.75E-11 | 6.28E+04 | 6.19E+01 | 2.58E-04 | 1.07E-06 |
| 7088 | 1.46E-08 | 2.80E-11 | 5.93E+04 | 7.07E+01 | 8.71E-04 | 1.29E-06 |
| MS-1805 | 2.54E-08 | 5.37E-11 | 5.72E+04 | 9.32E+01 | 1.45E-03 | 1.95E-06 |

NB: No Binding

### Table S2. X-ray data collection and refinement statistics for MAM01 Fab

| | MAM01 + Junctional region | MAM01 + Minor repeat region | MAM01 + Short major repeat region | MAM01 + Long major repeat region |
|---|---|---|---|---|
| **Data collection** | | | | |
| Beamline | SSRL 12-1 | SSRL 12-1 | SSRL 12-1 | SSRL 12-1 |
| Wavelength (Å) | 0.97946 | 0.97946 | 0.97946 | 0.97946 |
| Resolution (Å) | 30.00-2.71 (2.82-2.71)[a] | 30.00-1.84 (1.90-1.84)[a] | 30.00-1.54 (1.58-1.54)[a] | 30.00-1.48 (1.51-1.48)[a] |
| Space group | $P2_12_12_1$ | $P2_1$ | $P2_12_12_1$ | $P2_12_12_1$ |
| Unit cell a, b, c (Å) | 45.42, 72.81, 136.75 | 43.46, 66.34, 83.99 | 46.17, 71.55, 135.59 | 47.04, 71.53, 136.78 |
| α, β, γ (°) | 90, 90, 90 | 90, 96.1, 90 | 90, 90, 90 | 90, 90, 90 |
| Unique reflections | 13,394 (641) [a] | 37,764 (1967)[a] | 66,759 (3281)[a] | 77,121 (3808) [a] |
| Redundancy | 5.2 (5.6) [a] | 3.0 (3.1)[a] | 6.2 (6.3)[a] | 5.8 (6.5) [a] |
| Completeness (%) | 92.9 (93.5)[a] | 91.6 (93.3)[a] | 99.2 (98.5)[a] | 98.9 (99.4)[a] |
| Mean I/sigma($\sigma_I$) | 25.1 (4.5)[a] | 16.5 (1.7)[a] | 33.8 (3.7)[a] | 45.1 (4.5)[a] |
| $R_{sym}$ (%)[b] | 10.5 (44.9)[a] | 9.7 (65.7)[a] | 9.1 (57.0)[a] | 7.1 (50.6)[a] |
| $R_{pim}$ (%)[b] | 5.4 (19.9)[a] | 6.6 (42.7)[a] | 4.1 (22.4)[a] | 3.3 (21.6)[a] |
| $CC_{1/2}$ (%)[c] | 99.1 (95.3)[a] | 98.8 (73.2)[a] | 99.8 (99.4)[a] | 99.9 (90.3)[a] |
| **Refinement statistics** | | | | |
| Resolution (Å) | 29.43-2.71 | 28.56-1.84 | 29.44-1.54 | 29.77-1.48 |
| Reflections (work) | 11,963 | 37,700 | 66,533 | 76,892 |
| Reflections (test) | 1,198 | 1,999 | 1,999 | 1,998 |
| Rcryst[d] / Rfree[e] (%) | 26.0/31.6 | 19.4/23.3 | 17.8/20.3 | 19.3/20.8 |
| **Number of atoms** | | | | |
| Fab | 3,295 | 3,307 | 3,316 | 3,322 |
| Peptide | 93 | 108 | 84 | 104 |
| Water | 3 | 172 | 264 | 272 |
| **Average B-value (Å$^2$)** | | | | |
| Fab | 52 | 26 | 21 | 24 |
| Peptide | 52 | 27 | 14 | 20 |
| Water | 50 | 30 | 28 | 30 |
| Wilson B (Å$^2$) | 42 | 20 | 16 | 18 |
| **RMSD from ideal geometry** | | | | |
| Bond angle (°) | 0.66 | 0.97 | 1.05 | 1.15 |
| Bond length (Å) | 0.003 | 0.007 | 0.009 | 0.009 |
| **Ramachandran statistics**[f] | | | | |
| Favored (%) | 94.88 | 98.01 | 98.23 | 98.02 |
| Outliers (%) | 0.22 | 0.00 | 0.00 | 0.00 |
| **PDB Code** | 9NL1 | 9NZF | 9NL0 | 9NKZ |

[a] Numbers in parentheses refer to the highest resolution shell.
[b] Rsym = Σhkl Σi | Ihkl,i - | / Σhkl Σi Ihkl,i and Rpim = Σhkl (1/(n-1))1/2 Σi | Ihkl,i - | / Σhkl Σi Ihkl,i, where Ihkl,i is the scaled intensity of the ith measurement of reflection h, k, l, is the average intensity for that reflection, and n is the redundancy.
[c] CC1/2 = Pearson correlation coefficient between two random half datasets.
[d] Rcryst = Σhkl | Fo - Fc | / Σhkl | Fo | x 100, where Fo and Fc are the observed and calculated structure factors, respectively.
[e] Rfree was calculated as for Rcryst, but on a test set comprising 5% of the data excluded from refinement.
[f] From MolProbity[3].

### Table S3. X-ray data collection and refinement statistics for 7088 and MS-1805 Fabs

| | 7088 +N-term junctional region | 7088 + Short major repeat | MS-1805 +N-term junctional region | MS-1805 + Short major repeat |
|---|---|---|---|---|
| **Data collection** | | | | |
| Beamline | NSLS-II AMX | NSLS-II AMX | NSLS-II AMX | NSLS-II AMX |
| Wavelength (Å) | 0.92010 | 0.92010 | 0.92010 | 0.92010 |
| Resolution (Å) | 30.00-3.14 (3.14-3.20)[a] | 30.00-2.00 (2.14-2.00)[a] | 30.00-1.74 (1.78-1.74)[a] | 30.00-2.25 (2.29-2.25)[a] |
| Space group | $P2_12_12_1$ | $P4_32_12$ | $P2_1$ | $P2_1$ |
| Unit cell a, b, c (Å) | 72.00, 80.70, 99.38 | 72.11, 72.11, 195.79 | 42.33, 76.74, 74.45 | 50.54, 73.85, 60.75 |
| α, β, γ (°) | 90, 90, 90 | 90, 90, 90 | 90, 88.4, 90 | 90, 99.8, 90 |
| Unique reflections | 9,584 (464)[a] | 35,786 (1134)[a] | 45,572 (2226)[a] | 20,906 (1027)[a] |
| Redundancy | 12.8 (13.2)[a] | 11.5 (8.6)[a] | 3.2 (3.0)[a] | 6.7 (5.1) [a] |
| Completeness (%) | 94.2 (77.5)[a] | 99.6 (98.9)[a] | 93.1 (87.7)[a] | 98.9 (94.3)[a] |
| Mean I/sigma($\sigma_I$) | 11.5 (1.2)[a] | 61 (2.4)[a] | 21.40 (1.9)[a] | 11.5 (1.7)[a] |
| $R_{sym}$ (%)[b] | 19.5 (104)[a] | 10.9 (7.8)[a] | 6.9 (47.8)[a] | 12.7 (42.4)[a] |
| $R_{pim}$ (%)[b] | 5.6 (29.7)[a] | 3.4 (28.0)[a] | 4.5 (31.2)[a] | 5.3 (20.6)[a] |
| $CC_{1/2}$ (%)[c] | 99.9 (78.7)[a] | 98.1 (79.0)[a] | 99.5 (83.8)[a] | 98.2 (80.7)[a] |
| **Refinement statistics** | | | | |
| Resolution (Å) | 29.15-3.14 | 26.93-2.00 | 26.72-1.74 | 27.84-2.25 |
| Reflections (work) | 9,958 | 35,760 | 45,520 | 20,749 |
| Reflections (test) | 998 | 1,998 | 1,971 | 1,999 |
| Rcryst[d] / Rfree[e] (%) | 23.0/26.6 | 22.5/26.9 | 19.7/22.8 | 18.8/23.9 |
| **Number of atoms** | | | | |
| Fab | 3,294 | 3,265 | 3,296 | 3,302 |
| Peptide | 73 | 61 | 73 | 61 |
| Water | 0 | 19 | 163 | 78 |
| **Average B-value (Å²)** | | | | |
| Fab | 94 | 56 | 28 | 25 |
| Peptide | 130 | 50 | 23 | 27 |
| Water | 0 | 46 | 29 | 26 |
| Wilson B (Å²) | 79 | 46 | 20 | 23 |
| **RMSD from ideal geometry** | | | | |
| Bond angle (°) | 0.49 | 1.29 | 1.32 | 0.83 |
| Bond length (Å) | 0.002 | 0.012 | 0.014 | 0.006 |
| **Ramachandran statistics**[f] | | | | |
| Favored (%) | 94.38 | 95.04 | 98.38 | 97.45 |
| Outliers (%) | 0.23 | 0.24 | 0.00 | 0.23 |
| **PDB Code** | 9DSU | 9DSS | 9DST | 9DSR |

[a] Numbers in parentheses refer to the highest resolution shell.
[b] Rsym = Σhkl Σi | Ihkl,i - | / Σhkl Σi Ihkl,i and Rpim = Σhkl (1/(n-1))1/2 Σi | Ihkl,i - | / Σhkl Σi Ihkl,i, where Ihkl,i is the scaled intensity of the ith measurement of reflection h, k, l, is the average intensity for that reflection, and n is the redundancy.
[c] CC1/2 = Pearson correlation coefficient between two random half datasets.
[d] Rcryst = Σhkl | Fo - Fc | / Σhkl | Fo | x 100, where Fo and Fc are the observed and calculated structure factors, respectively.
[e] Rfree was calculated as for Rcryst, but on a test set comprising 5% of the data excluded from refinement.
[f] From MolProbity[3].

### Table S4. X-ray data collection and refinement statistics for unliganded Fabs

| | 224 unliganded | MAM01 unliganded | MS-1805 unliganded |
|---|---|---|---|
| **Data collection** | | | |
| Beamline | NSLS-II FMX | SSRL 12-1 | NSLS-II AMX |
| Wavelength (Å) | 0.97931 | 0.97946 | 0.92010 |
| Resolution (Å) | 30.00-1.70 (1.73-1.70)[a] | 30.00-2.08 (2.14-2.08)[a] | 30.00-1.77 (1.82-1.77)[a] |
| Space group | $P2_12_12_1$ | $P2_12_12_1$ | $P2_1$ |
| Unit cell a, b, c (Å) | 56.86, 72.83, 127.95 | 52.77, 69.79, 115.05 | 40.57, 145.4, 76.73 |
| α, β, γ (°) | 90, 90, 90 | 90, 90, 90 | 90, 91.4, 90 |
| Unique reflections | 60,033 (2821)[a] | 24,638 (1282)[a] | 85,659 (4239)[a] |
| Redundancy | 10.9 (7.8)[a] | 5.3 (65.6)[a] | 6.3 (5.0)[a] |
| Completeness (%) | 97.6 (83.9)[a] | 93.6 (94.6)[a] | 98.1 (90.9)[a] |
| Mean I/sigma($\sigma_I$) | 25.2 (0.7)[a] | 33.2 (5.32)[a] | 6.3 (0.9)[a] |
| $R_{sym}$ (%)[b] | 14.7 (87.9)[a] | 9.0 (43.0)[a] | 19.5 (71.0)[a] |
| $R_{pim}$ (%)[b] | 4.5 (31.5)[a] | 4.7 (20.2)[a] | 8.4 (34.8)[a] |
| $CC_{1/2}$ (%)[c] | 99.4 (65.2)[a] | 99.7 (95.1)[a] | 98.1 (41.8)[a] |
| **Refinement statistics** | | | |
| Resolution (Å) | 26.48-1.70 | 29.11-2.08 | 26.31-1.77 |
| Reflections (work) | 59,090 | 24,567 | 84,405 |
| Reflections (test) | 2000 | 1999 | 2010 |
| Rcryst[d] / Rfree[e] (%) | 20.0/23.1 | 23.1/27.0 | 20.4/23.1 2 |
| **Number of atoms** | | | |
| Fab | 3,267 | 3,244 | 6,577 |
| Water | 207 | 29 | 389 |
| **Average B-value (Å$^2$)** | | | |
| Fab | 33 | 45 | 18 |
| Water | 36 | 39 | 22 |
| Wilson B (Å$^2$) | 26 | 38 | 12 |
| **RMSD from ideal geometry** | | | |
| Bond angle (°) | 1.01 | 0.73 | 0.82 |
| Bond length (Å) | 0.009 | 0.005 | 0.005 |
| **Ramachandran statistics**[f] | | | |
| Favored (%) | 97.68 | 98.59 | 98.22 |
| Outliers (%) | 0.00 | 0.00 | 0.00 |
| **PDB Code** | 9NOB | 9NKY | 9NOA |

[a] Numbers in parentheses refer to the highest resolution shell.
[b] Rsym = Σhkl Σi | Ihkl,i - | / Σhkl Σi Ihkl,i and Rpim = Σhkl (1/(n-1))1/2 Σi | Ihkl,i - | / Σhkl Σi Ihkl,i, where Ihkl,i is the scaled intensity of the ith measurement of reflection h, k, l, is the average intensity for that reflection, and n is the redundancy.
[c] CC1/2 = Pearson correlation coefficient between two random half datasets.
[d] Rcryst = Σhkl | Fo - Fc | / Σhkl | Fo | x 100, where Fo and Fc are the observed and calculated structure factors, respectively.
[e] Rfree was calculated as for Rcryst, but on a test set comprising 5% of the data excluded from refinement.
[f] From MolProbity[3].
